# PfClpC Is An Essential Clp Chaperone Required For Plastid Integrity And Clp Protease Stability In *Plasmodium falciparum*

**DOI:** 10.1101/080408

**Authors:** Anat Florentin, David W Cobb, Jillian D Fishburn, Michael J Cipriano, Paul S Kim, Manuel A Fierro, Boris Striepen, Vasant Muralidharan

## Abstract

The deadly malaria parasite, *Plasmodium falciparum*, contains a non-photosynthetic plastid known as the apicoplast, that functions to produce essential metabolites. Little is known about its biology or regulation, but drugs that target the apicoplast are clinically effective. Using phylogenetic analysis, we identified a putative complex of clp (caseinolytic protease) genes. We genetically targeted members of this complex and generated conditional mutants of the PfClpC chaperone and PfClpP protease and found that they co-localize in the apicoplast. Conditional inhibition of the PfClpC chaperone resulted in growth arrest and apicoplast loss, and was rescued by addition of the essential apicoplast-derived metabolite, IPP. Using a double conditional-mutant parasite line, we discovered that the chaperone activity is required to stabilize the active protease, revealing functional interactions. These data demonstrate the essential function of PfClpC in maintaining apicoplast integrity and its role in regulating the proteolytic activity of the Clp complex.

## Introduction

Malaria is a devastating human disease caused by obligate intracellular parasites of the genus *Plasmodium*. This disease results in nearly 450,000 deaths each year, which are mostly caused by one species, *Plasmodium falciparum* (World Health Organization, 2015). The parasite has gained resistance to all clinically available antimalarial drugs, generating an urgent need to identify new drugs and potential novel drug targets (Hovlid and Winzeler, 2016; Wells et al., 2015). The parasite cell is remarkably complex with two organelles that carry their own genetic material, the mitochondrion and a unique algal endosymbiont known as the apicoplast (McFadden, 1996). The apicoplast harbors essential metabolic pathways that are required for parasite growth and survival (van Dooren and Striepen, 2013). Importantly, drugs that target cellular processes in the apicoplast are clinically effective (Erica L Dahl, 2007; Fichera and Roos, 1997; Goodman et al., 2007). Therefore, understanding the function, structure and biogenesis of the apicoplast provides a rich vein of antimalarial drug targets. One potential class of such targets are the <u>c</u>aseino<u>l</u>ytic <u>p</u>rotease (Clp) family of proteins that act as key regulators of the biology of bacterial cells, the evolutionary ancestors of the apicoplast. In bacteria and plant chloroplasts, Clp proteins play vital roles in cell/ organelle division, segregation, protein homeostasis and protein transport (Frees et al., 2014; Nishimura and van Wijk, 2015). Typically, they form a regulated proteolytic complex in which a Clp protease is paired with a Clp chaperone that has a AAA+ ATPase domain (also known as the Hsp100 family of chaperones) such as ClpC or ClpA (Olivares et al., 2016). There are several putative *clp* genes encoded in the *P. falciparum* genome which have been localized to the apicoplast (Bakkouri et al., 2010), and previous studies using recombinant proteins described structural features of the Clp protease and its inactive subunit (Bakkouri et al., 2013; Rathore et al., 2010). Currently, nothing is known about the apicoplast-localized Clp chaperones, and their roles *in vivo* as well as the interactions between the apicoplast Clp proteins remain poorly understood due to the challenging genetics of *P. falciparum*.

In this study, we generated conditional mutants of the *P. falciparum* apicoplast-targeted *pfclpc* and *pfclpp* genes and found that they localize to the apicoplast. Conditional inhibition of the PfClpC chaperone resulted in growth arrest, morphological defects, and apicoplast loss. In a series of cellular assays, we showed that PfClpC is required for apicoplast sorting into daughter cells. Addition of IPP, an essential apicoplast-derived metabolite, rescued growth, indicating that the only essential function of PfClpC is linked to the apicoplast. Using a double conditional-mutant *pfclpc; pfclpp* parasite line, we discovered that PfClpC activity is required to stabilize PfClpP in its active-protease form, revealing a functional interaction between the two. Our work demonstrates the essential function of PfClpC in maintaining apicoplast integrity and segregation and its role in regulating the proteolytic activity of the Clp complex.

## Results

### Phylogenetic analysis of *Plasmodium* Clp Proteins

In order to identify Clp proteins that potentially form a proteolytic complex and compare them to other organisms, we performed phylogenetic analyses of putative Clp proteases and chaperones (Figure 1A,B). Our analyses show that PfClpP is the only ClpP protease in the *P. falciparum* plastid, unlike other apicomplexans, which encode several copies (Figure 1A). PfClpP is closely related to ClpP of plants and cyanobacteria, in agreement with the origin of the apicoplast in the secondary endosymbiosis of a red alga. Importantly, ClpP homologs are present throughout eubacterial kingdom, including many pathogenic species, which has led to the development of potent specific inhibitors (Böttcher and Sieber, 2008; Brötz-Oesterhelt and Sass, 2014; Conlon et al., 2013; Gersch et al., 2015; Hackl et al., 2015). The genome of *P. falciparum* also encodes PfClpR, a ClpR homolog, which is related to ClpP but lacks catalytic residues. PfClpR is grouped with ClpR orthologues from cyanobacteria and plants, indicating that it did not result from a recent gene duplication event (Figure 1A). Phylogenetic analysis of *Plasmodium* Clp chaperones assigns each one of them to a different subfamily, such as ClpB1, ClpB2, ClpC and ClpM (Figure 1B). Among those, there is a single ClpC orthologue, PfClpC, which is the only Clp chaperone with a predicted ClpP binding motif. It falls into a clade together with plants, cyanobacteria, and *Haemosporidia*, a sub-class of apicomplexans. Other apicomplexans such as the closely related parasite *Toxoplasma gondii*, do not encode ClpC homologs, but contain ClpM and multiple duplications of ClpB1 and ClpB2. This suggests variations in the Clp complexes within apicomplexans, and that PfClpC may have a *Plasmodium* specific function.

**Figure 1:**
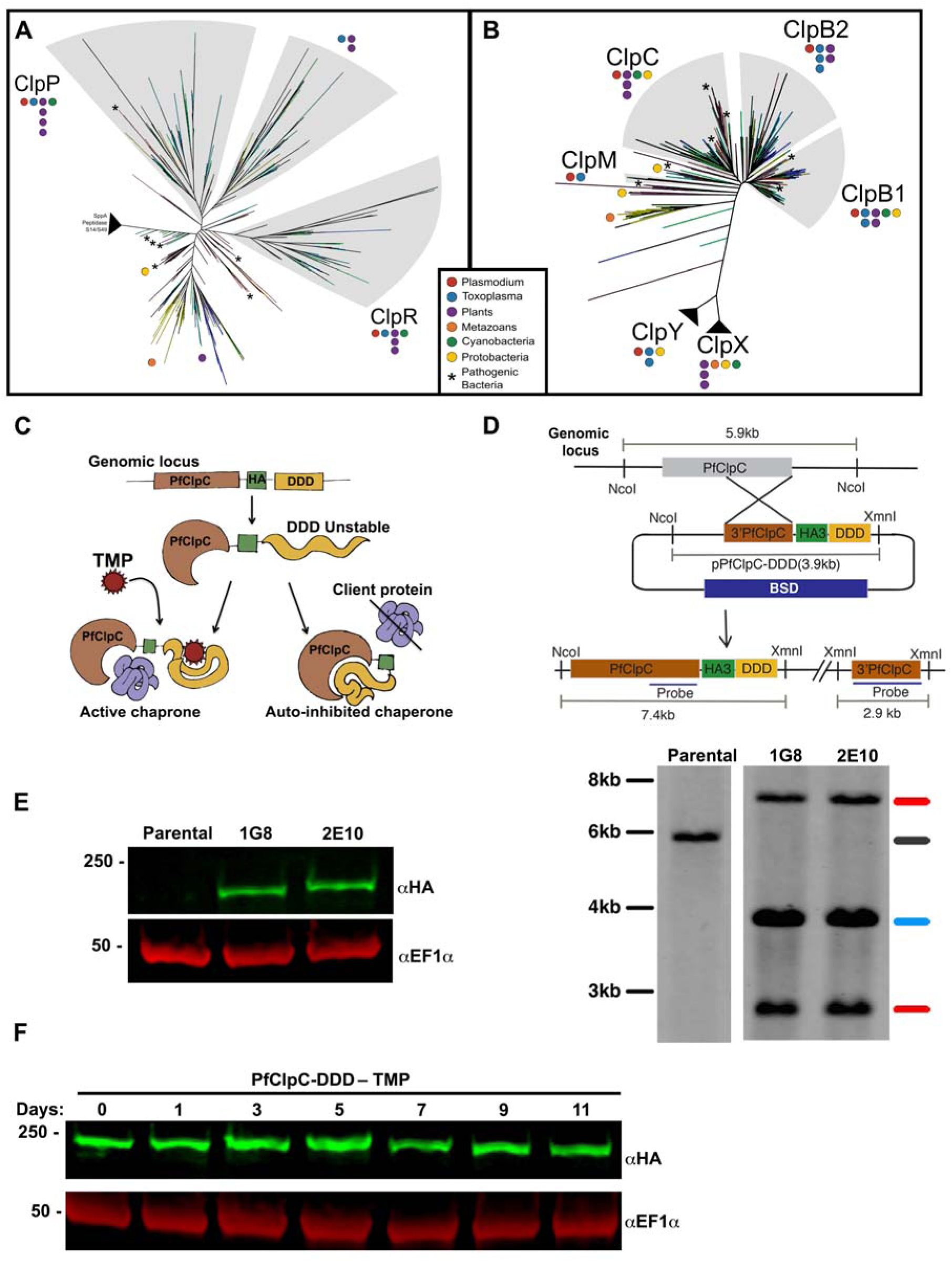
Phylogenetic analysis and generation of PfClpC-DDD transgenic parasites. a) Phylogenetic analysis of the ClpP proteases. The unrooted maximum likelihood tree was created using RaxML and represents the best scoring tree. Shaded areas define certain sub-families of Clp proteases. Colored dots represent ClpP homologs in different species/ taxa. More than one dot with the same color within a given subfamily denote gene duplication. For representation purposes dots are grouped together and are not located near individual branches to simplify the image. Asterisks denote medically important pathogenic bacteria with ClpP homologs including *Mycobacterium tuberculosis, Staphylococcus aureus, Streptococcal bacteria, Neisseria meningitides, Salmonella, Escherichia coli* and *Bacillus anthracis.* For the full tree and complete alignment sequences see supplemental information. b) Phylogenetic analysis of Clp chaperones. The unrooted maximum likelihood tree was created using FastTree2. Shaded areas define subfamilies of Clp chaperones. Colored dots represent Clp chaperones in different species/ taxa. More than one dot with the same color within a given subfamily denote gene duplication. *Plasmodium* species encode only one ClpC, and it is the only Clp chaperone with a predicted ClpP binding motif. PfClpC falls into a clade together with plants, cyanobacteria, and *Haemosporidia*, a sub-class of apicomplexan. For representation purposes dots are grouped together and are not located near individual branches to simplify the image. Asterisks denote medically important pathogenic bacteria with homologs of Clp chaperones including *Mycobacterium tuberculosis, Staphylococcus aureus, Streptococcal bacteria, Neisseria meningitides, Salmonella, Escherichia coli* and *Bacillus anthracis.* For the full tree and complete alignment sequences see supplemental information. c) Mechanism of PfClpC conditional inhibition. The *pfclpc* locus was modified to contain a triple hemagglutinin (HA) tag and a DHFR-based destabilization domain (DDD). A small molecule ligand, Trimethoprim (TMP), is used to stabilize the DDD and maintain normal chaperone function. Upon TMP removal the chaperone binds the DDD intramolecularly and cannot interact with client proteins, thus inhibiting normal chaperone activity. d) Single crossover homologous recombination enables the integration of the ClpC-DDD plasmid into the 3’ end of the *pfclpc* gene (upper panel). Southern blot analysis of PfClpC-DDD genomic DNA (lower panel) isolated from parasite lines indicated above the lanes. The genomic DNA was digested with NcoI and XmnI. Bands expected from integration of the plasmid into the 3’ end of the *pfclpc* gene were observed in two clones (1G8 and 2E10), isolated from two independent transfections (Red). A plasmid band was observed in the clones (blue), suggesting that a plasmid concatamer integrated into the gene. A single band indicative of the parental allele was observed for the parental strain (black) and it was absent in the integrant clones. e) Western blot of parasite lysates from parental line and two independent PfClpC-DDD clones (1G8 and 2E10) probed with antibodies against HA (green) and EF1α (loading control, red). The protein marker sizes that co-migrated with the probed protein are shown on the left. f) TMP was removed from PfClpC-DDD parasites and parasite lysates were isolated every 24 or 48 hours over 11 days. PfClpC and EF1α were visualized on Western blots using antibodies against HA (PfClpC-DDD, green) and EF1α (loading control, red). The protein marker sizes that co-migrated with the probed protein are shown on the left. One representative experiment out of four is shown (two for each clone).

### Generating conditional mutants for PfClpC and PfClpP

Based on this analysis, we took a genetic approach to dissect the biological roles of the *Plasmodium* putative Clp complex, consisting of PfClpP, PfClpR, and PfClpC. To study the function of the chaperone PfClpC, we inserted a DHFR-based destabilizing domain (DDD) into the *pfclpc* genomic locus, a technique used for conditional auto-inhibition of chaperone function (Beck et al., 2014; Muralidharan et al., 2012). In the chaperone-DDD fusion protein, the unfolded DDD binds the chaperone intra-molecularly, thereby excluding client proteins and inhibiting normal chaperone function (Figure 1C). A small molecule ligand, trimethoprim (TMP) is used to stabilize and refold the DDD, releasing the chaperone to resume its normal function (Figure 1C). Using a single-crossover homologous recombination approach, we tagged the C-terminus of the *pfclpc* gene with a triple-HA tag and the DDD to produce the PfClpC-DDD parasite line (Figure 1D). Two clones from independent transfections were isolated (1G8 and 2E10) and integration was confirmed by Southern blot analysis (Figure 1D). Using an anti-HA antibody, we detected expression of the PfClpC-DDD fusion protein at the expected molecular size in tagged clones but not in the parental line (Figure 1E). As predicted by the chaperone auto-inhibition model, TMP removal did not lead to PfClpC degradation (Figure 1F).

To study PfClpP and PfClpR, we used a translational inhibition technique, involving the *glmS* ribozyme. In this method, a *glmS* sequence is inserted into the genomic locus before the 3’UTR of the gene, after the stop codon (Prommana et al., 2013). Addition of the small molecule glucosamine (GlcN) results in its conversion inside the cell to glucosamine 6-phosphate which activates the *glmS* ribozyme. The now-active ribozyme cleaves itself from the mRNA, leading to transcript degradation. Using CRISPR/Cas9, we appended a triple-V5 tag to the C-terminus of PfClpP and PfClpR followed by the *glmS* sequence (Figure 2A). We successfully modified the *pfclpp* genomic locus and obtained two independent parasites lines, PfClpP-glmS1 and PfClpP-glmS2, in which correct integration was verified by genomic PCR (Figure 2B). Using an anti-V5 antibody, we confirmed expression, and observed that PfClpP is proteolytically processed and appears as a full-length inactive-zymogen (I), apicoplast-localized zymogen without the transit peptide (II) and an active protease after cleavage of the pro-domain (III) (Figure 2C)(Bakkouri et al., 2010; Rathore et al., 2010). Upon addition of GlcN, PfClpP protein levels were reduced by 60% (Figure 2D). The same strategy was used for tagging *pfclpr* locus, however integration was inefficient, and multiple attempts to clone mutants from mixed populations via limiting dilution failed, suggesting that PfClpR is essential for asexual growth (Figure S1).

**Figure 2:**
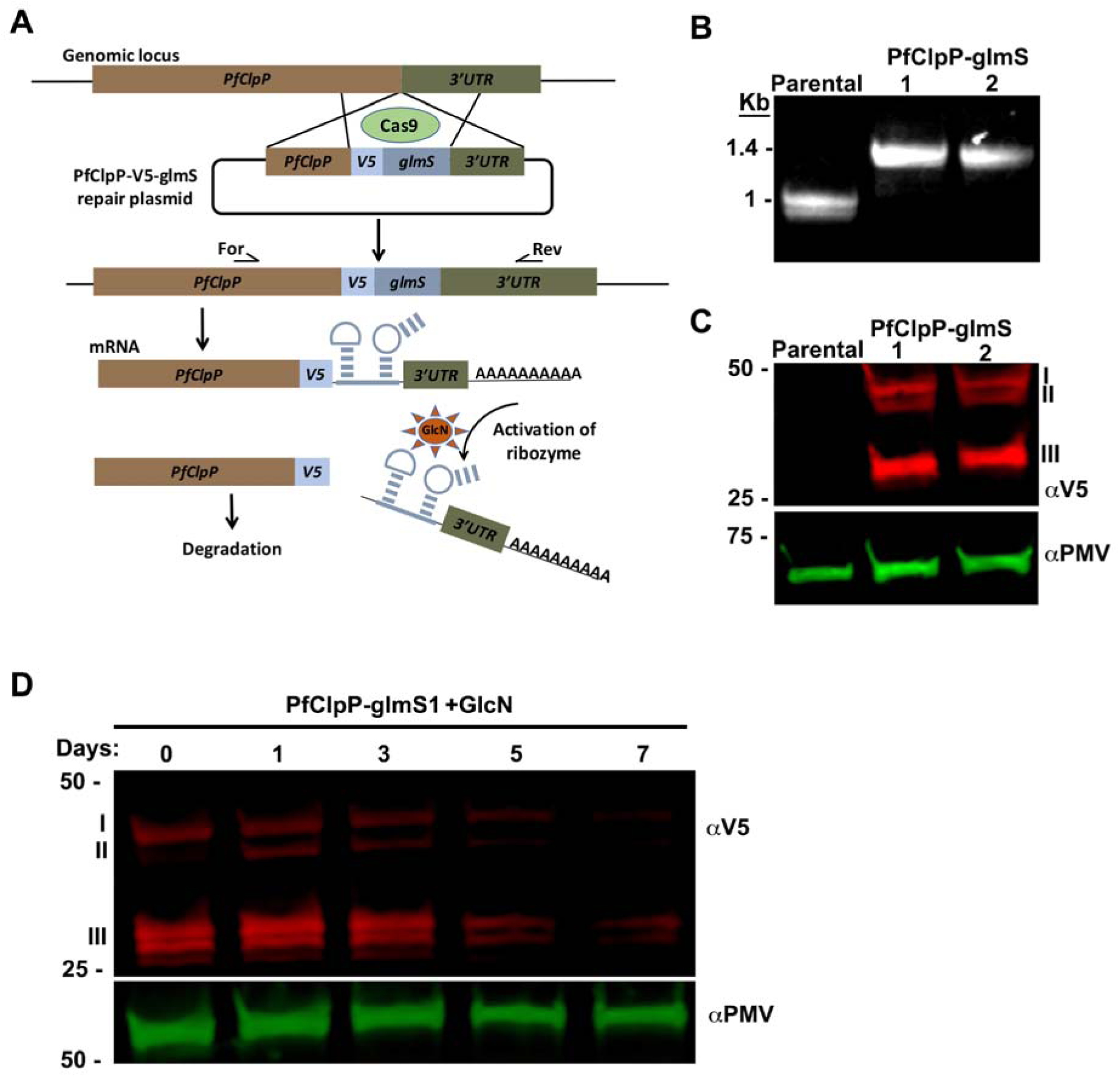
Generating PfClpP-glmS transgenic parasites. a. Diagram showing integration of the *glmS*-ribozyme sequence into the *pfclpp* locus, and the mechanism of GlcN-induced degradation of mRNA. Cas9 introduces a double-stranded break at the beginning of the 3’ UTR of the *pfclpp* locus. The repair plasmid provides homology regions for double-crossover homologous recombination, introducing a 3xV5 tag and the glmS-ribozyme sequence. Addition of the small molecule glucosamine (GlcN) results in its conversion inside the cell into glucosamine 6-phosphate which activates the *glmS* ribozyme. The now-active ribozyme cleaves itself from the mRNA, leading to transcript degradation. b. PCR test confirming *V5-glmS* integration at the *pfclpp* locus. DNA was purified from transfected parasites and primers 9 and 8 (see under methods) were used to amplify the region between the 3’ end of the ORF and 3’UTR of *pfclpp*, as illustrated in the diagram. A shift of 400 bp corresponds to the integration of the 3xV5 tag and the glmSribozyme sequence. c. Western blot analysis of parasite lysates from parental line and two independent transfections (PfClpP-glmS1 and PfClpP-glmS2) probed with antibodies against V5 (red) and Plasmepsin V (PMV, loading control, green). PfClpP is proteolyticly processed and the different bands correspond to the full-length inactive-zymogen (I), apicoplast-localized, transit peptide-cleaved (II) and active protease, pro-domain removed (III). The protein marker sizes that co-migrated with the probed protein are shown on the left. d. GlcN was added to PfClpP-glmS parasites and parasite lysates were isolated every 24 or 48 hours over 7 days. PfClpP and EF1α were visualized on a Western blot using antibodies against V5 (PfClpP-glmS, red) and Plasmepsin V (PMV, loading control, green). A 60% reduction in the total protein levels is observed. The protein marker sizes that co-migrated with the probed protein are shown on the left. One representative experiment out of four is shown (two for each transfection).

To detect the sub-cellular localization of PfClpC and PfClpP we used Immunofluorescence microscopy involving staining for known apicoplast resident proteins. We were able to determine the apicoplast localization of PfClpC by staining with anti-HA and anti-acyl carrier protein (ACP, an apicoplast marker (Waller et al., 1998)) and PfClpP by staining with anti-V5 and anti-Cpn60 (homolog of the *T. gondii* apicoplast chaperone Cpn60 (Agrawal et al., 2009)) (Figure 3A).

**Figure 3:**
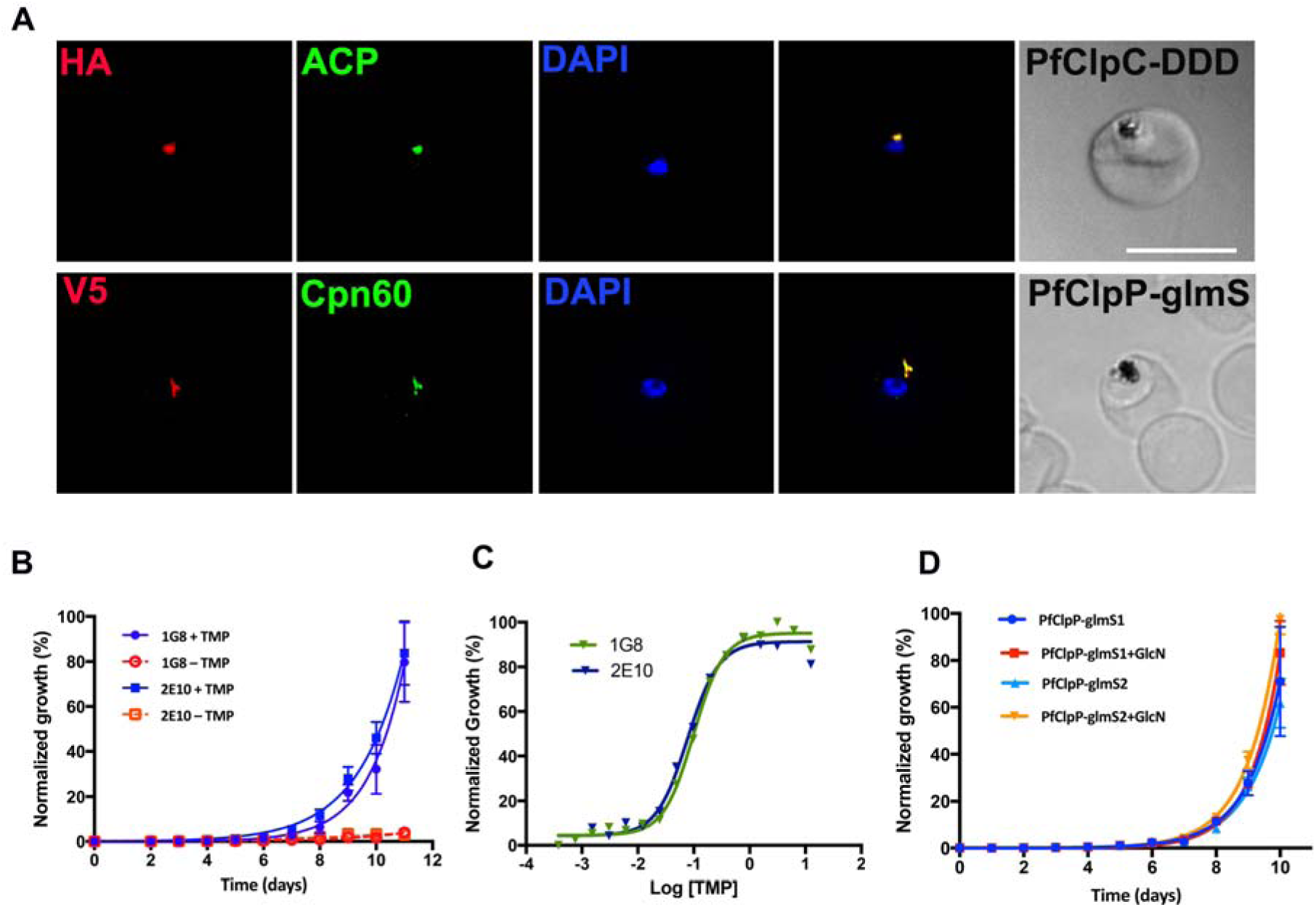
PfClpC Activity is Essential for Parasite Growth within the Red Blood Cell. A. Immunofluorescence imaging of fixed unsynchronized PfClpC-DDD parasites (1G8) stained with antibodies against HA (red), ACP (green) and DAPI (blue) (upper panel) and PfClpP-glmS parasites (PfClpP-glmS1) stained with antibodies against V5 (red), Cpn60 (green) and DAPI (blue) (lower panel). Z-stack images were deconvolved and projected as a combined single image. Scale bar, 5μm. B. Asynchronous PfClpC-DDD clones, 1G8 and 2E10, were grown with or without 10μM TMP and parasitemia was monitored every 24 hours over 12 days via flow cytometry. During the course of the experiment parasite cultures were sub-cultured to maintain parasitemia between 1-5% and data were calculated using the actual parasitemia multiplied by the dilution factors of each individual culture. 100% of relative parasitemia represents the highest value of calculated parasitemia. Growth inhibition is observed after 7-8 days post TMP removal, corresponding to roughly 3-4 asexual cycles. Data are fit to an exponential growth equation and are represented as mean ± S.E.M. One representative experiment out of three is shown. C. Asynchronous PfClpC-DDD parasites were incubated for 11 days without TMP, and on day 12 were seeded in a 96 well plate with varying concentrations of TMP. Parasitemia was measured after 5 days using flow cytometry showing an EC50 of 80nM. Data are fit to a dose-response equation and are represented as mean ± S.E.M. One representative experiment out of four is shown. D. Asynchronous PfClpP-glmS parasites were grown with or without 5mM GlcN and parasitemia was monitored every 24 hours over 10 days via flow cytometry. During the course of the experiment parasite cultures were sub-cultured to maintain parasitemia between 1-5% and data were calculated using the actual parasitemia multiplied by the dilution factors of each individual culture. 100% of relative parasitemia represents the highest value of calculated parasitemia. Data are fit to an exponential growth equation and are represented as mean ± S.E.M.

### PfClpC is essential for asexual growth of parasites

To test the requirement of PfClpC for asexual replication, we removed the stabilizing ligand (TMP) and monitored the growth of PfClpC-DDD parasites. We did not see a strong effect initially, and since *P. falciparum* asexual life cycle takes 48 hours, we concluded that that these parasites develop normally for the first two or three growth cycles. However, a significant growth defect was observed seven days after TMP withdrawal and eventually lead to a severe growth arrest in these parasites (Figure 3B). We found that inhibition of PfClpC activity relies on TMP in a dose-dependent manner with an EC50 of 80nM (Figure 3C). Addition of GlcN to PfClpP-glmS parasites did not affect asexual growth (Figure 3D), most likely due to insufficient reduction in PfClpP levels (Figure 2D). While we could not determine the effect of PfClpP inhibition, we concluded that PfClpC activity is essential for parasite survival and asexual growth within human red blood cells.

### PfClpC inhibition leads to a reduced replication rate within the red blood cells

Further characterization of PfClpC mutants revealed that they developed normally during the early stages of rings and trophozoites (Figure S2A). Late schizont stages (≥ 6 nuclei), however, developed aberrant morphology, including irregular cellular shape, empty vacuoles and fewer nuclei, suggesting that they are nonviable (Figure 4A). These morphologically abnormal parasites appeared on the 3^rd^ replication cycle and their fraction increased over time (Figure 4B). Analysis of synchronized late-stage parasites over several replication cycles using flow cytometry revealed that instead of a typical single peak, they had a wider distribution, suggesting variation in DNA content (Figure 4C). To test the replication efficiency of these mixed populations of parasites, we monitored the rate of schizonts to rings conversion. We found that TMP removal resulted in a significant decrease in the numbers of ring-stage parasites that were formed in each successive generation (Figure 4D). This reduced replication rate accounts for the observed growth inhibition as well as the increase in the numbers of morphologically abnormal parasites with each replication cycle.

**Figure 4:**
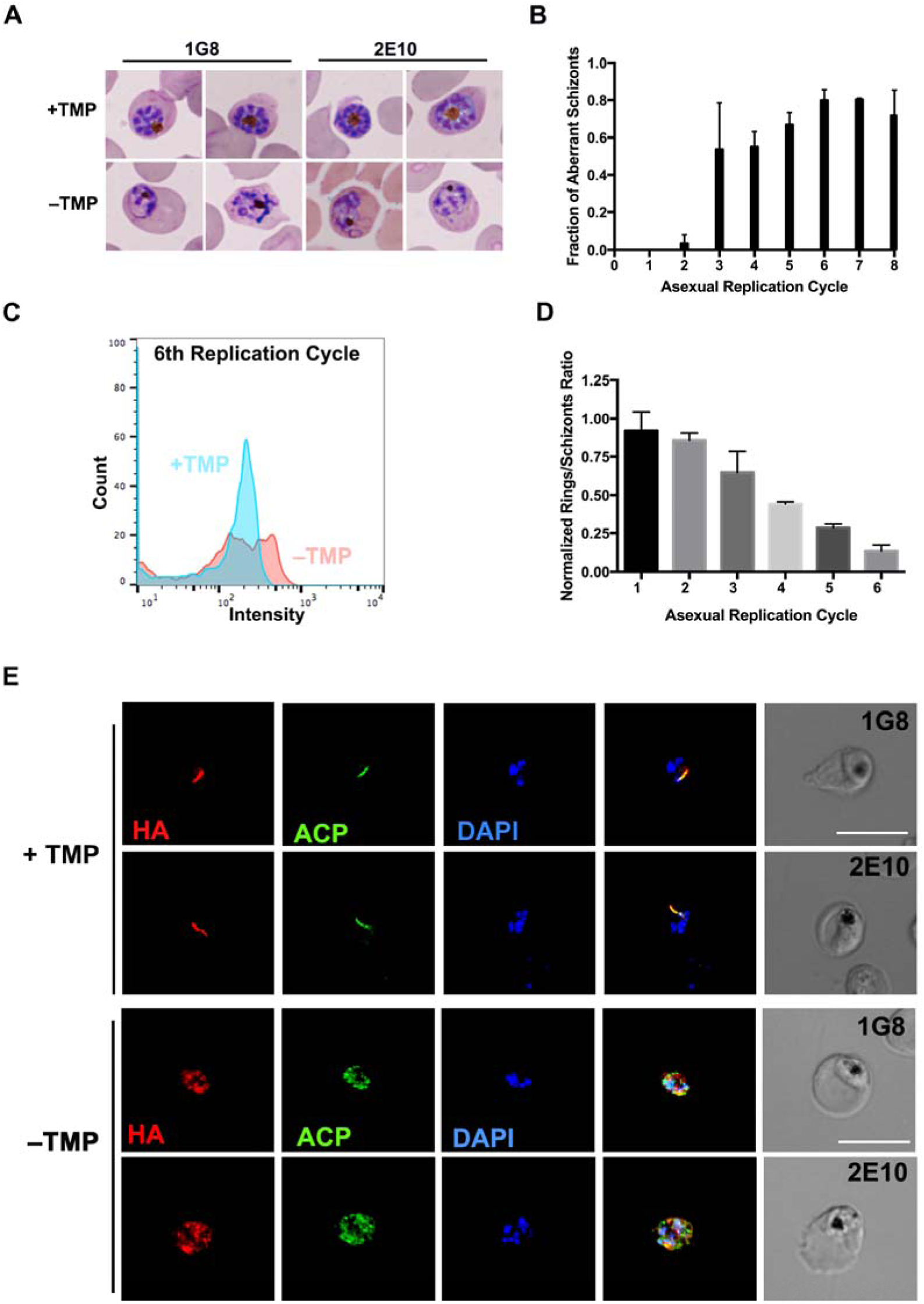
PfClpC inhibition reduces replication rate and apicoplast integrity. a) Hema 3 stained thin blood smears of PfClpC-DDD parasites that were grown for 10 days without TMP. Both clones of PfClpC-DDD parasites are shown, as indicated above the images. b) TMP was removed from synchronized PfClpC-DDD parasites and thin blood smears of late stage schizonts were stained and analyzed using light microscopy. Parasites were counted, and the fraction of defective cells (as seen in 4A) was calculated out of the total late-stage population (≥6 nuclei). One asexual replication cycle is roughly 48 hours. Data are shown from one representative experiment with clone 2E10. Two experiments (with technical triplicates) were performed with each clone. c) TMP was removed from synchronized PfClpC-DDD parasites and the DNA of the parasites was stained using Acridine Orange and was analyzed by flow cytometry during the 6^th^ replication cycle. One asexual replication cycle is roughly 48 hours. One representative image is shown (out of four experiments, two with each clone). d) TMP was removed from synchronized PfClpC-DDD parasites and the numbers of rings and late schizont stages was determined by flow cytometry. The ratio of rings to schizonts was calculated using the number of rings arising from schizonts in the previous generation. Cultures were re-synchronized at the beginning of each replication cycle using sorbitol. One asexual replication cycle is roughly 48 hours. Data were normalized to the ring: schizont ratio in the presence of TMP. Data are shown for 1G8 clone. The experiment was performed twice for each clone (with technical triplicates in each experiment). e) PfClpC-DDD parasites were incubated for 10 days without TMP and then fixed and stained with antibodies against HA (red), ACP (green) and DAPI (blue). Both clones of PfClpC-DDD parasites are shown as indicated. Images from left to right are anti-HA, anti-ACP, DAPI, fluorescence merge and phase. Z-stack images were deconvolved and projected as a combined single image. Scale bar, 5μm.

### PfClpC activity is required for apicoplast integrity

To test a possible effect of PfClpC inhibition on the apicoplast, we removed TMP and performed immunofluorescence microscopy assays and observed a loss of canonical apicoplast morphology. Instead, we detected PfClpC, as well as the apicoplast resident protein ACP, in a punctate, vesicle-like pattern, suggesting that apicoplast integrity was compromised (Figure 4E). Apicoplast targeting of nuclear-encoded proteins is mediated through an N-terminal transit peptide that is cleaved upon localization to the apicoplast (Foth et al., 2003; Waller, 2000). In line with the loss of apicoplast localization, a second higher band for PfClpC appeared upon TMP removal, indicative of a cytoplasmic fraction of this protein in which the transit peptide was not cleaved (Figure S2B).

### Chemical rescue of PfClpC-DDD parasites using IPP

The only essential function of the apicoplast during the blood stages is the biosynthesis of isopentenyl-pyrophosphate (IPP), the precursor for all isoprenoids, through the non-mevalonate (or MEP) pathway (Yeh and DeRisi, 2011). To test the effect of IPP on PfClpC inhibition, we removed TMP and added IPP to PfClpC-DDD parasites. We were able to observe complete restoration of normal growth as well as typical cellular morphology (Figure 5A,B). Immunofluorescence microscopy revealed, however, that IPP-treated PfClpC-DDD parasites survived in the absence of a functional apicoplast and retained the vesicle-like structures containing PfClpC and ACP (Figure 5C). To further investigate the fate of the apicoplast in PfClpC-DDD parasites, we performed quantitative Real Time PCR (qRT-PCR) experiments, comparing the nuclear, plastid and mitochondrial genomes over subsequent replication cycles without TMP. This qRT-PCR analysis revealed a significant reduction in the plastid genome, whereas the mitochondrial genome was unaffected (Figure 5D). This indicates that in addition to a functional damage to the apicoplast, there was an actual loss of the plastid genome. Overall, we concluded that the only essential activity of PfClpC is linked to apicoplast function.

**Figure 5:**
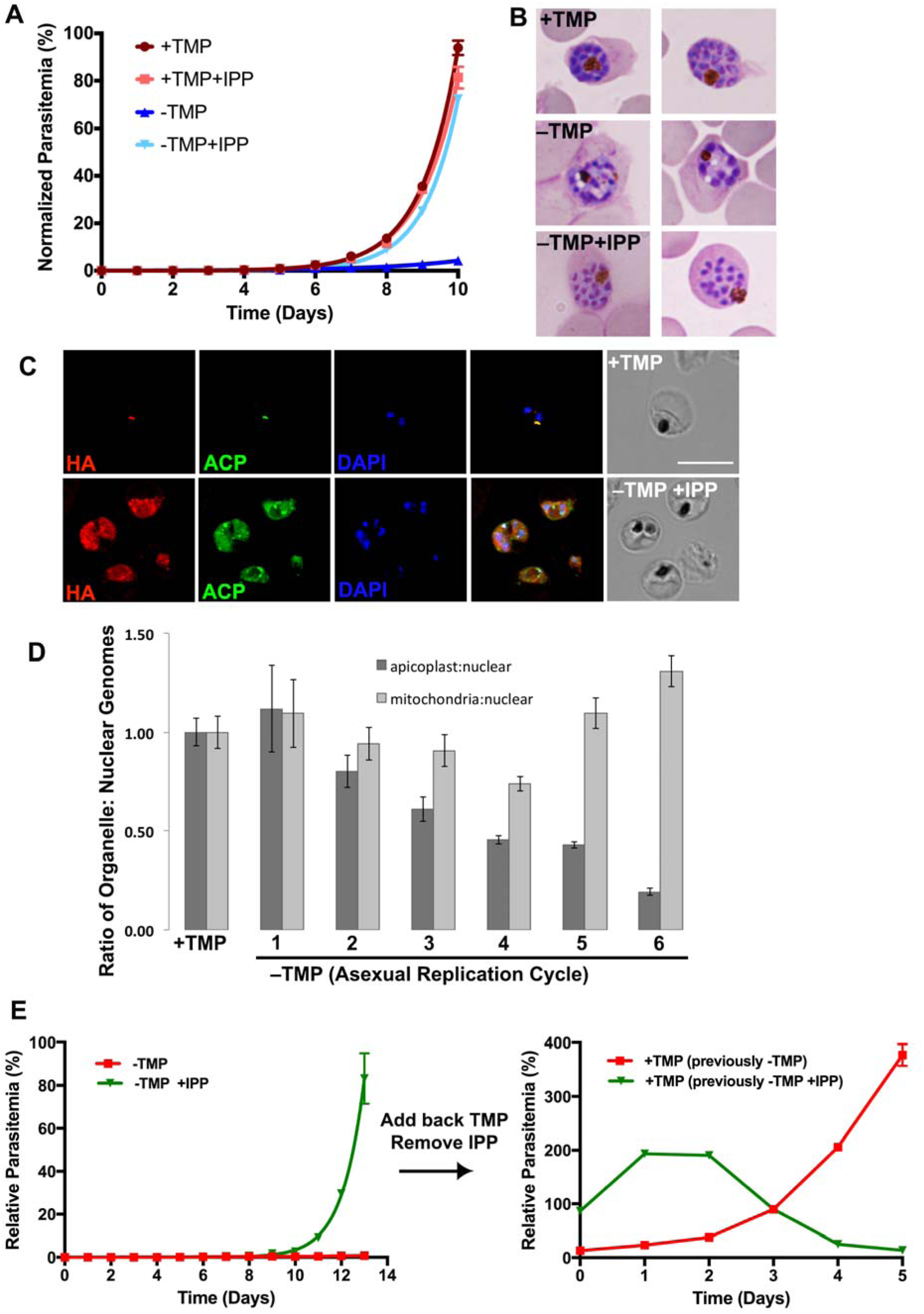
Chemical Rescue of PfClpC Auto-inhibition using IPP. a) PfClpC-DDD parasites were grown for 10 days without TMP and supplemented with isopentenyl pyrophosphate (IPP). Parasitemia was measured using flow cytometry. During the course of the experiment parasite cultures were sub-cultured to maintain parasitemia between 1-5% and data were calculated using the actual parasitemia multiplied by the dilution factors of each individual culture. 100% of relative parasitemia represents the highest value of calculated parasitemia at the end of the experiment. Data are fit to an exponential growth equation and are represented as mean ± S.E.M. One of two (one for each clone) representative experiments is shown. Graph denotes data collected for 1G8 clone. b) Hema 3 stained thin blood smears of PfClpC-DDD parasites (1G8 clone) grown for 10 days with TMP (upper), without TMP (middle) or without TMP and supplemented with IPP (bottom). Two representative images for each condition are shown. c) PfClpC-DDD parasites (1G8 clone) grown for 10 days without TMP and supplemented with IPP. These PfClpC-DDD parasites were fixed and stained with antibodies against HA (red) and ACP (green). Images from left to right are anti-HA, anti-ACP, DAPI, fluorescence merge and phase. Z-stack images were deconvolved and projected as a combined single image. Scale bar, 5μm. d) Synchronized PfClpC-DDD parasites were grown in the absence of TMP and presence of IPP and DNA samples were taken at the beginning of each replication cycle for quantitative Real Time PCR analysis. Apicoplast: nuclear genome ratio was calculated for each replication cycle. Mitochondria: nuclear genome ratio served as a control. One asexual replication cycle is roughly 48 hours. Genome ratios were normalized to parasites grown in the presence of TMP. Data are represented as mean ± S.E.M. one representative experiment is shown (out of four, two for each clone). e) PfClpC tagged parasites were grown for 14 days without TMP (red line), or without TMP and supplemented with IPP (green line). During the course of the experiment parasite cultures were sub-cultured to maintain parasitemia between 1-5% and data were calculated using the actual parasitemia multiplied by the dilution factors of each individual culture. 100% of relative parasitemia represents the highest value of calculated parasitemia at the end of the experiment. Data are fit to an exponential growth equation and are represented as mean ± S.E.M. On day 14 IPP was removed from the media and TMP was added back to all parasite cultures and parasitemia was measured using flow cytometry. 100% of relative parasitemia represents the highest parasitemia value at the beginning of the experiment. Data are represented as mean ± S.E.M. One representative experiment out of two for each clone is shown.

### An apicoplast sorting defect in PfClpC-DDD parasites

The reduced replication rate upon loss of PfClpC function could be indicative of a problem in apicoplast segregation into daughter cells. To test this, we aimed to visualize apicoplast presence or absence in early stage parasites (≤ 5 hours post-invasion). We used imaging flow cytometry to analyze these parasites and found that PfClpC was present in all parasites (Figure S3A and Table S1). Higher resolution microscopy confirmed that these early stage parasites contained PfClpC and it was present in vesicle-like punctate pattern rather than in the canonical apicoplast structure (Figure S3B). This suggests that either young parasites inherit these vesicles from their mother cell, or that these vesicles represent very early *de novo* synthesis of these proteins. Either way, nuclear-encoded apicoplast proteins are expressed and present in the cell, even in the earliest stages, regardless of apicoplast presence or absence.

We therefore employed a functional assay to differentiate between parasites containing an apicoplast and parasites that have lost it, relying on the fact that the plastid must be inherited and cannot be synthesized *de novo*. We removed TMP and allowed PfClpC-DDD parasites to grow in the presence or absence of IPP for two weeks. As expected, PfClpC-DDD parasites grown without TMP and supplemented with IPP grew normally, whereas parasites incubated without TMP and without IPP were unable to grow and were undetectable for several days (Figure 5E, left). On day 14 we removed IPP and relieved PfClpC inhibition by adding back TMP, and monitored the growth of the parasites. Upon adding back TMP to the media, the parasites that were grown without IPP recovered and resumed normal growth, indicating that a small fraction of inhibited parasites still possessed a functional apicoplast (Figure 5E, right). Conversely, parasites that were grown with IPP started dying 48 hours after removing IPP and adding back TMP (Figure 5E, right). These parasites lost their apicoplast but used IPP to survive, and therefore died despite restoration of PfClpC activity. Overall, these data indicate that PfClpC-DDD mutants lose the apicoplast from most parasites, with the exception of a small population that retains the plastid, suggesting a sorting defect.

### PfClpC is required to stabilize PfClpP active protease form

Since PfClpC is the only Clp chaperone in the apicoplast with a ClpP binding motif (Figure 1B) (Bakkouri et al., 2010; Kim et al., 2001), we were interested in possible interactions between the chaperone and the other Clp complex subunits. Multiple attempts to immunoprecipitate PfClpC or PfClpP in their native-state from parasite-cell lysates have failed, probably due to the compact nature of the Clp complex that renders the epitope tags inaccessible to the antibodies. We also explored the effect of a previously described bacterial ClpP inhibitor, U1-lactone, that was shown to inhibit PfClpP *in vitro* (Rathore et al., 2010). We observed, however, that inhibition of parasite growth by U1-lactone could not be rescued by IPP, indicating that *in vivo* it is not specific for PfClpP and has at least one target outside of the apicoplast (Figure S4). Therefore, to explore functional interactions between PfClpC and PfClpP, we generated *pfclpc; pfclpp* double-conditional mutants. In these parasites, PfClpC is tagged with HA and is controlled by TMP removal and PfClpP is tagged with V5 and is controlled by addition of GlcN (Figure 6A,B). These parasites responded to PfClpC inhibition but not to partial reduction of PfClpP, similar to the single mutants (Figure 6C). Interestingly, PfClpC inhibition affected PfClpP localization, and it appeared in vesicle-like structures together with PfClpC (Figure 6D). This was accompanied with accumulation of the of the full-length PfClpP-zymogen, as seen by the appearance of the band with the highest apparent molecular weight on an SDS-PAGE (I) (Figure 6E). These observations were made seven days or more after TMP removal, and were consistent with the growth arrest and apicoplast loss in these parasites. Unexpectedly, the second processing event of PfClpP, i.e. removal of the pro-domain to produce the active protease, was also inhibited upon chaperone inactivation, but with much faster kinetics (Figure 6E). One day after TMP removal we observed significant reduction in the active protease form (III), and accumulation of the inactive zymogen (II) (Figure 6E). Analysis at shorter time points revealed a rapid decrease in the active form of PfClpP within 4 hours after TMP removal (Figure 6F). This rapid effect suggests that PfClpC activity is required to stabilize PfClpP in its active form.

**Figure 6:**
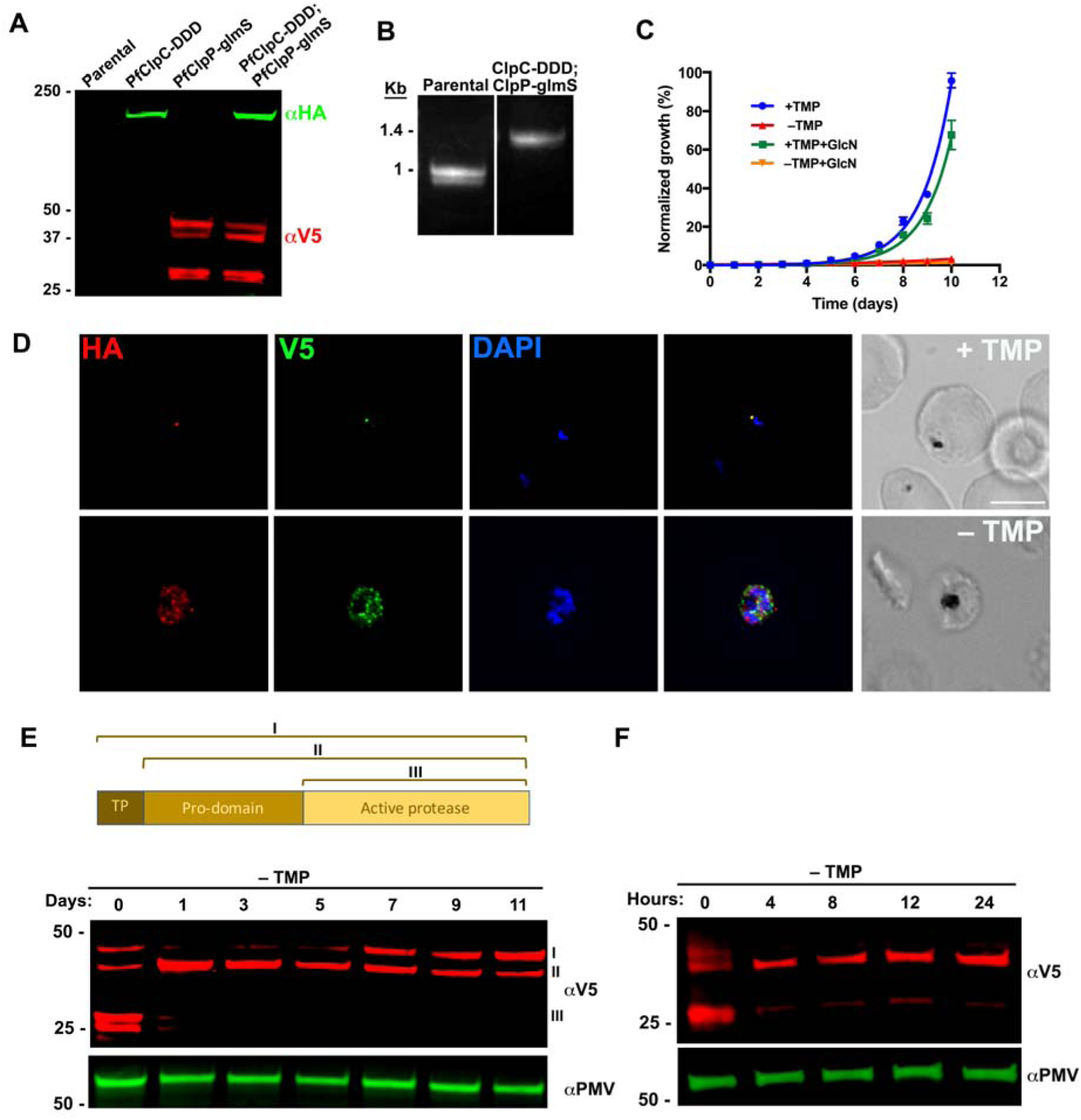
PfClpC is required for PfClpP localization and stability. a) Western blot of parasite lysates from parental line (3D7), PfClpC-DDD, PfClpP-glmS, and PfClpC-DDD; PfClpP-glmS (double mutant) parasites probed with antibodies against HA (green) and V5 (red). The protein marker sizes that co-migrated with the probed protein are shown on the left. b) PCR test confirming *V5-glmS* integration at the *pfclpp* locus in a PfClpC-DDD background. DNA was purified from transfected parasites and primers 9 and 8 were used to amplify the region between the C-terminus and 3’UTR of *pfclpp*, as illustrated in Figure 3a. c) Asynchronous PfClpC-DDD; PfClpP-glmS parasites were grown with or without 10μM TMP/ 5mM GlcN and parasitemia was monitored every 24 hours over 12 days via flow cytometry. During the course of the experiment parasite cultures were sub-cultured to maintain parasitemia between 1-5% and data were calculated using the actual parasitemia multiplied by the dilution factors of each individual culture. 100% of relative parasitemia represents the highest value of calculated parasitemia. Data are fit to an exponential growth equation and are represented as mean ± S.E.M. d) PfClpC-DDD; PfClpP-glmS parasites were incubated for 10 days without TMP and then fixed and stained with antibodies against HA (red), V5 (green) and DAPI (blue). Z-stack images were deconvolved and projected as a combined single image. Scale bar, 5μM. e) Upper panel: Schematic representation of the full-length PfClpP protein with the Transit Peptide (I). Removal of the Transit Peptide (TP) occurs upon localization to the apicoplast (II). Self-cleavage of the pro-domain occurs upon assembly into an active complex (III). Lower panel: PfClpC-DDD; PfClpP-glmS (double mutant) parasites were incubated without TMP for 11 days and lysates were isolated every 24 or 48 hours and probed with antibodies against V5 (red) and PMV (loading control, green). The protein marker sizes that co-migrated with the probed protein are shown on the left. f) PfClpC-DDD; PfClpP-glmS (double mutant) parasites were incubated without TMP for 24 hours and lysates were isolated every 4-12 hours and probed with antibodies against V5 (red) and PMV (loading control, green). The protein marker sizes that co-migrated with the probed protein are shown on the left.

## Discussion

The deadly malaria parasite, *Plasmodium falciparum*, is a eukaryotic pathogen and as such, it shares conserved basic biology with its human host. It is therefore both challenging and essential, in the search for potential drug targets, to identify key components that are absent or significantly different from the human host.

One such class of potential candidates is the apicoplast-associated prokaryotic Clp family of chaperones and proteases. In the bacterial ancestors, as well as in other organellar descendants such as the mitochondria and chloroplast, these proteins serve a variety of basic molecular functions ranging from protein degradation, transport across membranes, protein folding, cell division, stress response and pathogenicity (Frees et al., 2014). Little is known, however, about the physiological roles of the apicoplast-resident Clp proteins in the biology of *Plasmodium falciparum*.

In this study, we performed a thorough phylogenetic analysis, and identified the Clp proteins that potentially form a proteolytic complex in the apicoplast of *P. falciparum*. These include the protease PfClpP, the inactive subunit PfClpR, and PfClpC, a AAA+ chaperone with a unique ClpP binding motif (Figure 1). This analysis revealed homology to the Clp proteins of several pathogenic bacteria as well as to those that are in plants chloroplasts, but not necessarily to other apicomplexan parasites such as *Toxoplasma gondii*. This suggests that parasites have adopted and use this machinery in different ways, pointing, in particular in the case of the PfClpC chaperone, to a *Plasmodium* specific function.

Taking a genetic approach, we were able to demonstrate the localization of PfClpP and PfClpC to the apicoplast, and showed that the chaperone activity is essential to parasite viability and asexual replication (Figure 3). In a series of cellular assays, we were able to show that it is required for apicoplast integrity. Inhibition of PfClpC resulted in the loss of distinct apicoplast morphology and in the presence of vesicular-like structures (Figure 4). Several studies reported the appearance of such structures when the apicoplast integrity is compromised, for example with the use of certain antibiotics (Gisselberg et al., 2013; Yeh and DeRisi, 2011). This was interpreted as stalled vesicular transport from the ER that cannot dock to the apicoplast membrane due to loss of the organelle (van Dooren and Striepen, 2013). Moreover, chemical rescue using IPP restored PfClpC-DDD growth and cellular morphology despite the absence of the apicoplast as could be seen by microscopy and qRT-PCR (Figure 5). It has been shown that isoprenoid biosynthesis is the only essential metabolic function of the apicoplast, and supplementing IPP can replace a non-functional apicoplast in living parasites (Yeh and DeRisi, 2011). We therefore concluded that the only essential role of PfClpC is linked to apicoplast function. Inhibition of essential apicoplast metabolic pathways with drugs like Fosmidomycin, kills parasites immediately and does not lead to the loss of the organelle (Bowman et al., 2014). Conversely, inhibition of apicoplast translation or replication with drugs like Doxycycline, allows the parasites to complete one replication cycle, proceed through the next, and die only during the second schizogony (Fichera and Roos, 1997; Yeh and DeRisi, 2011). Similar to the effect of drugs that inhibit apicoplast replication, PfClpC mutants developed normally during the early stages of rings and trophozoites but late schizont stages developed aberrant morphology (Figure 4). These non-viable parasites did not manifest uniformly at the end of the second cycle but appeared on the 3^rd^ replication cycle, and their fraction increased over time (Figure 4B). A possible explanation for the delayed growth arrest, as well as the gradual increase in abnormal parasites, is that PfClpC inhibition interferes with the segregation or division of functional apicoplast into daughter merozoites. As a consequence, a mixed population of viable and non-viable daughter cells is forming after each cycle, diluting overtime the viable parasites in the total culture. Indeed, we observed a significant decrease in the rate of schizont to ring conversion in each successive generation, clarifying the delayed growth inhibition, and suggesting a defect in apicoplast sorting (Figure 4).

To support that, we performed functional analysis, by inhibiting and then reactivating the chaperone function. This was achieved by removing TMP from the cultures for two weeks, reduce parasite numbers below detection, and then adding back TMP to reactivate the chaperone in any possible remaining parasites (Figure 5E). In the event of a uniform functional damage to all parasites in the culture, PfClpC re-activation would not lead to viable parasites, as the apicoplast must be inherited and its *de novo* synthesis is impossible. Nonetheless, we observed that re-addition of TMP could restore parasites growth, indicating the presence of a small yet undetectable sub-population of parasites that contains a functional apicoplast. This observation further supports a sorting defect rather than a general apicoplast dysfunction in the entire parasite population. Interestingly, TMP addition had the opposite effect on parasites that were rescued with IPP (Figure 5E). These parasites started dying 48 hours after removing IPP despite addition of TMP, indicating that re-activation of PfClpC was not enough to sustain viability in a population of parasites that permanently lost the apicoplast.

In addition, we describe here a mechanism by which PfClpC regulates the activity of the Clp complex. We found that PfClpC activity is necessary and regulates the processing of PfClpP protease in several ways (Figure 6). Chaperone inhibition resulted in PfClpP miss-localization and reduced the cleavage of its transit peptide 7 days or more after TMP removal. These observations are correlated and consistent with the kinetics of apicoplast loss in PfClpC mutants (Figure 3, 4 and 5). But it also affected the active form of PfClpP with much faster kinetics. We detected a significant reduction in PfClpP active-protease form (in which the transit peptide and the pro-domain have been removed) 4 hours after TMP removal, suggesting degradation of the complex subunits when the chaperone is inhibited (Figure 6F). In this model, the activity of the Clp chaperone is required to stabilize the proteolytic Clp complex, representing a novel mode of regulation, which may be relevant to bacterial and chloroplasts systems.

Finally, the roles that bacterial Clp proteins play in cell division, stress response and pathogenicity (Frees et al., 2014), have placed them at the center of several drug discovery programs (Böttcher and Sieber, 2008; Brötz-Oesterhelt and Sass, 2014; Conlon et al., 2013; Gersch et al., 2015; Hackl et al., 2015). Our data demonstrate that targeting the *P. falciparum* plastid-localized Clp proteins is a viable strategy for antimalarial drug development and future work will allow us to repurpose highly active antibacterial compounds as effective anti-malarial agents.

## Experimental Procedures

### Multiple sequence alignments and phylogenetic analyses

Candidate proteins were chosen based on the presence of conserved Pfam domains (Punta et al., 2012), CLP_protease for the Clp protease family and ClpB_D2-small for the Clp chaperone family. Representative genomes were searched using hmmscan, then candidate full length proteins were aligned with clustalo (Sievers and Higgins, 2014) using the Pfam generated hmm as a guide, then the alignment was trimmed with trimal (Capella-Gutiérrez et al., 2009) with the following settings “-st 0.00001 -gt 0.01”. Exact duplicate alignments were temporarily removed prior to the creation of trees and added back as 0.00001 distance sister taxa.

Maximum likelihood trees of candidate proteins were creating using FastTree2 (Price et al., 2010) with options “-spr 4 -mlacc 2 -slownni -gamma” or RaXML (Stamatakis, 2014) using the best estimate protein model selection using the script ProteinModelSelection.pl from RaXML and RaXML options “-f a -x 12345 -p 12345 -N autoMRE“. Rogue taxa (those taxa with uncertain positions in phylogenetic trees) were identified with RagNaRok (Aberer et al., 2013) and excluded from trees. Trees were then visualized and interrogated with Archeopetryx (Han and Zmasek, 2009).

Complete trees and full sequence alignemnts can be found in supplemental information.

### Plasmids construction

Genomic DNAs were isolated from *P. falciparum* using the QIAamp DNA blood kit (QIAGEN). Constructs utilized in this study were confirmed by sequencing. PCR products were inserted into the respective plasmids using the In-Fusion cloning system (Clonetech) or using SLIC (Sequence and Ligation Independent Cloning). Briefly, insert and cut vector were mixed with a T4 DNA polymerase and incubated for 2.5 minutes at room temperature, followed by 10 minutes incubation on ice and then transformed into bacteria. All restriction enzymes used for this study were purchased from New England Biolabs. All oligonucleotides and detailed cloning procedures can be found under Supplemental experimental procedures

### Cell culture and transfections

Parasites were cultured in RPMI medium supplemented with Albumax I (Gibco) and transfected as described earlier (Drew et al., 2008; Russo et al., 2009). To generate PfClpC-DDD parasites, pPfClpC-HADB was transfected in duplicates into 3D7-derived parental strain PM1KO which contains a hDHFR expression cassette conferring resistance to TMP (Liu et al., 2005). Selection, drug cycling and cloning were performed as described (Muralidharan et al., 2012) in the presence of 10 μM of TMP (Sigma). Integration was detected after one round of drug cycling with blasticidin (Sigma). Two clones from 2 independent transfections, 1G8 and 2E10, were isolated via limiting dilutions and used for subsequent experiments.

For generation of PfClpP-glmS and PfClpR-glmS parasites, a mix of two plasmids was transfected in duplicates into 3D7 parasites. The plasmids mix contained pUF1-Cas9-guide (Spillman et al., 2017) which contains the *DHOD* resistance gene, and pV5-glmS-PfClpP or pV5-glmS-PfClpR, which are marker-free. To generate the double-mutant parasites, PfClpC-DDD; PfClpP-glmS, the same plasmid mix was transfected into PfClpC-DDD parasites. Drug pressure was applied 48 hours post transfection, using 1 μM DSM1 (Ganesan et al., 2011), selecting only for Cas9 expression. DSM1 was removed from the culturing media once parasites were detected in the culture, usually around 3 weeks post transfection.

### Growth assays

For asynchronous growth assays, parasites were washed twice and incubated without TMP (for PfClpC-DDD parasites) or with 5 mM GlcN (Sigma) (for PfClpP*-glmS* parasites). Throughout the course of the experiment parasites were sub-cultured to maintain the parasitemia between 1-5% and parasitemia was monitored every 24 hours via flow cytometry. Relative parasitemia at each time point was back calculated based on actual parasitemia multiplied by the relevant dilution factors. Parasitemia in the presence of TMP (PfClpC-DDD) or in the absence of GlcN (PfClpP-glmS) at the end of each experiment was set as the highest relative parasitemia and was used to normalize parasites growth. Data were fit to exponential growth equations using Prism (GraphPad Software, Inc.)

For IPP rescue, media was supplemented with 200 μM of IPP (Isoprenoids LC) in PBS. To test for the ClpP inhibitor, media was supplemented with or without 60 μM of U1-lactone and 200 μM of IPP.

To generate an EC50 curve for TMP, asynchronous PfClpC-DDD parasites were incubated for 11 days without TMP, and on day 12 were seeded in a 96 well plate with varying concentrations of TMP. Parasitemia was measured after 5 days using flow cytometry. Data were fit to a dose-response equation using Prism (GraphPad Software, Inc.).

### Replication rate analysis

To determine replication rate (rings: schizonts ratio) of PfClpC-DDD parasites, TMP was removed from percoll-isolated schizonts-stage parasites and parasites were allowed to egress and reinvade fresh RBCs. Parasitemia was monitored by flow cytometry and microscopy. The ratio of rings to schizonts was calculated using number of rings arising from schizonts in the previous generation. At the beginning of each replication cycle parasites were re-synchronized using Sorbitol, and sub-cultured when required. For each replication cycle, data were normalized to rings: schizonts ratio in the presence of TMP.

The fraction of morphologically aberrant schizonts was determined using Giemsa-stained thin blood smears of synchronized PfClpC-DDD parasites at the final stages of each replication cycle and the fraction of defective cells was calculated based on the total late schizont stage parasite counts.

### Southern blot

Southern blots were performed with genomic DNA isolated using the Qiagen Blood and Cell Culture kit. 10 μg of DNA isolated from PfClpC-DDD parasites was digested overnight with NcoI and XmnI (New England Biolabs) and integrants were screened using biotin-labeled probes against the 3’-end of the *pfclpc* ORF. Southern blot was performed as described earlier (Klemba et al., 2004). The probe was labeled using biotinylated Biotin-16-dUTP (Sigma). The biotinylated probe was detected on blots using IRDye 800CW Streptavidin conjugated dye (LICOR Biosciences) and was imaged, processed and analyzed using the Odyssey infrared imaging system software (LICOR Biosciences).

### Western blot

Western blots were performed as described previously (Muralidharan et al., 2011). Briefly, parasites were collected and host red blood cells were permeabilized selectively by treatment with ice-cold 0.04% saponin in PBS for 10 min, followed by a wash in ice-cold PBS. Cells were lysed using RIPA buffer, sonicated, and cleared by centrifugation at 4°C. The antibodies used in this study were rat monoclonal anti-HA, 3F10 (Roche, 1:3000), rabbit anti-V5, D3H8Q (Cell Signaling, 1:1000), mouse monoclonal anti-PMV (from D. Goldberg, 1:400), and rabbit polyclonal anti-EF1α (from D. Goldberg, 1:2000). The secondary antibodies that were used are IRDye 680CW goat anti-rabbit IgG and IRDye 800CW goat anti-mouse IgG (LICOR Biosciences, 1:20,000). The Western blot images were processed and analyzed using the Odyssey infrared imaging system software (LICOR Biosciences).

### Microscopy and image processing

For IFA cells were fixed using a mix of 4% Paraformaldehyde and 0.015% glutaraldehyde and permeabilized using 0.1% Triton-X100. Primary antibodies used are rat anti-HA clone 3F10 (Roche, 1:100), rabbit anti-V5, D3H8Q (Cell Signaling, 1:100), mouse anti-V5, TCM5 (eBioscience™, 1:100), rabbit anti-Cpn60 (1:1,000) and rabbit anti-ACP (from G. Mcfadden, 1:10,000). Secondary antibodies used are Alexa Fluor 488 and Alexa Fluor 546 (Life Technologies, 1:100). Cells were mounted on ProLong Diamond with DAPI (Invitrogen) and were imaged using DeltaVision II microscope system with an Olympus IX-71 inverted microscope using a 100X objective. All images were collected as Z-stack, were deconvolved using the DVII acquisition software SoftWorx and displayed as maximum intensity projection. Image processing, analysis and display were preformed using SoftWorx and Adobe Photoshop. Adjustments to brightness and contrast were made for display purposes. Thin blood smears were stained using Hema 3 stain set (PRTOCOL/ Fisher Diagnostics) and were imaged on a Nikon Eclipse E400 microscope.

### Flow cytometry

Aliquots of parasite cultures (5μl) were stained with 1.5 mg/ml Acridine Orange (Molecular Probes) in PBS. The fluorescence profiles of infected erythrocytes were measured by flow cytometry on a CyAn ADP (Beckman Coulter, Hialeah, Florida) and analyzed by FlowJo software (Treestar, Inc., Ashland, Oregon). The parasitemia data were fit to standard growth curve or dose–response using Prism (GraphPad Software, Inc.).

### Quantitative Real Time PCR

Synchronized ring stage parasites samples were collected at the beginning of each replication cycle and genomic DNA was isolated by saponin lysis to remove extracellular DNA. Genomic DNA was purified using QIAamp blood kits (Qiagen). Primers that amplify segments from genes encoded by nuclear or organelles genomes were designed using RealTime qPCR Assay Entry (IDT). The following primer sequences were used: *cht1* (nuclear): 21 + 22. *tufA* (apicoplast): 23 + 24. *cytb3* (mitochondria): 25 + 26. Reactions contained template DNA, 0.5 μM of gene specific primers, and IQ™ SYBR Green Supermix (BIORAD). Quantitative real-time PCR was carried out in triplicates and was performed at a 2-step reaction with 95°C denaturation and 56°C annealing and extension for 35 cycles on a CFX96 Real-Time System (BIORAD). Relative quantification of target genes was determined using Bio-Rad CFX manager 3.1 software. Standard curves for each primers set were obtained by using different dilutions of control gDNA isolated from parasites grown in the presence of TMP (20 to 0.2 ng) as template, and these standard curves were used to determine primers efficiency. For each replication cycle number, the organelle: nuclear genome ratio of the –TMP+IPP treated parasites was calculated relative to that of the +TMP control.

### Imaging flow cytometry

Synchronized PfClpC-DDD parasites incubated for 10 days without TMP and then were isolated on a percoll gradient following by a sorbitol treatment 5 hours later to obtain early rings (0-5 hours post invasion). Cells were fixed and stained with anti HA antibody as described above and nuclei were stained using DAPI from Amnis Intracellular staining kit (EMD MILIPORE). Data were collected on ImageStream X Mark II (EMD MILIPORE) and an automated collection of a statistically large number of cells (10,000) was performed. Data were analyzed using IDEAS software version 6.2.

## Acknowledgements

We thank Geoffrey McFadden for anti-ACP antibody and Dan Goldberg for pUF1-Cas9 plasmid, anti-PMV and anti EF1α antibodies; Stephan A. Sieber for providing the U1-lactone inhibitor. Drew Etheridge and Roberto Docampo for comments on the manuscript; Julie Nelson at the CTEGD Cytometry Shared Resource Laboratory for help with flow cytometry and analysis; and Muthugapatti Kandasamy at the Biomedical Microscopy Core at the University of Georgia for help with microscopy. We acknowledge the assistance of the Children’s Healthcare of Atlanta and Emory University Pediatric Flow Cytometry Core for imaging flow cytometry. This work was supported by grants from the March of Dimes Foundation (Basil O’Connor Starter Scholar Research Award), the US National Institutes of Health (R00AI099156 and R21AI128195), and UGA startup funds to V. M.

